# Rapid Surge of Reassortant A(H1N1) Influenza Viruses in Danish Swine and their Zoonotic Potential

**DOI:** 10.1101/2024.12.11.627926

**Authors:** Pia Ryt-Hansen, Sophie George, Charlotte Kristiane Hjulsager, Ramona Trebbien, Jesper Schak Krog, Marta Maria Ciucani, Sine Nygaard Langerhuus, Jennifer DeBeauchamp, Jeri Carol Crumpton, Taylor Hibler, Richard J. Webby, Lars Erik Larsen

## Abstract

In 2018, a single detection of a novel reassortant swine influenza A virus (swIAV) was made in Denmark. The hemagglutinin (HA) of the virus was from the H1N1 pandemic 2009 (H1N1pdm09) lineage and the neuraminidase (NA) from the H1N1 Eurasian avian-like swine lineage (H1N1av). By 2022, the novel reassortant virus (H1pdm09N1av) constituted 27 % of swIAVs identified through the Danish passive swIAV surveillance program. Sequencing detected two H1pdm09N1av genotypes; Genotype 1 contained an internal gene cassette of H1N1pdm09 origin, Genotype 2 differed by carrying an NS gene segment of H1N1av origin. The internal gene cassette of Genotype 2 became increasingly dominant, not only in the H1pdm09N1av population, but also in other Danish enzootic swIAV subtypes. Phylogenetic analysis of the HA genes from H1pdm09N1av viruses revealed a monophyletic source, a higher substitution rate compared to other H1N1pdm09 viruses and genetic differences with human seasonal and other swine adapted H1N1pdm09 viruses. Correspondingly, H1pdm09N1av viruses were antigenically distinct from human H1N1pdm09 vaccine viruses. Both H1pdm09N1av genotypes transmitted between ferrets by direct contact, but only Genotype 1 was capable of efficient aerosol transmission. The rapid spread of H1pdm09N1av viruses in Danish swine herds is concerning for swine and human health. Their zoonotic threat is highlighted by the limited pre-existing immunity observed in the human population, aerosol transmission in ferrets and the finding that the internal gene cassette of Genotype 2 was present in the first two zoonotic infections ever detected in Denmark.

## Introduction

During the last 15 years, two viral pandemics emerged from animal reservoirs, underlining the importance of viral surveillance in animal species in close contact with humans. Intensification of swine production systems over the past two decades has significantly increased the density of pigs in swine herds. Selective breeding of sows with larger litter sizes leads to enhanced mixing of immunological naïve piglets and age groups in many farms. These conditions complicate effective biosecurity and facilitates the continuous circulation of multiple enzootic swine influenza A viruses (swIAVs), which in turn increases the risk of reassortment (1–3).

An increasing number of swIAV reassortants have been reported globally since the introduction of the H1N1pdm09 virus in 2009 (4–8). Reassortant swIAVs not only pose a threat to swine herds, as novel swIAVs can escape herd immunity, increase transmission and cause severe disease (9–11), but also have unpredictable zoonotic potential. The last three human influenza pandemics were all caused by novel IAV reassortants (12). Simultaneously with reassortment events a constant genetic drift of swIAVs within the individual herds is seen and is further enhanced by intensive swine production that creates optimal conditions for continuous swIAV circulation. Genetic drift is driven by herd immunity stimulated by both swIAV infections and repeated mass vaccinations (13–15).

In Denmark, early reverse zoonotic transmission of H1N1pdm09 (1A.3.3.2) viruses led to the generation of H1pdm09N2 swIAV viruses and their internal gene cassette was found in most circulating swIAV strains. More recently, in 2018, a novel reassortant was detected that had a Eurasian avian-like (H1N1av) swIAV NA resulting in a novel virus termed “H1pdm09N1av” (16).

In 2021, the first two zoonotic IAV infections were registered in Denmark, both were caused by reassortant swIAVs. The first case was caused by a swine H1N1pdm09 virus (A/Denmark/1/2021(H1N1v)) (17) and the second was caused by a swine H1pdm09N1av virus (A/Denmark/36/2021(H1N1v)) (18); both had NS gene segments of H1N1av origin.

The aim of this study was to investigate the genetic nature of swIAV collected as part of the Danish swine surveillance program from 2019-2022 and to investigate the zoonotic risk of representative viruses.

## Materials and methods

### Samples, screening and lineage determination

Since 2010, Denmark has conducted a passive, prospective surveillance program for swIAV in Danish swine herds. Data obtained from the surveillance program during 2019-2022 were included in this study. The structure of the surveillance program, including laboratory analyses, have been previously described (16). In brief, Danish veterinarians submitted samples from pigs experiencing clinical signs of respiratory disease to one of three veterinary laboratories; Statens Serum Institut (SSI), Danish Technical University (DTU) and Veterinary Laboratory, Danish Agriculture & Food Council (L&F). The samples included nasal swabs, oral fluid and lung tissue. RNA was extracted from the samples using two different methods; the RNeasy mini kit automated on the QIAcube Connect (Qiagen, Germany) (DTU and SSI) or the MagNA Pure 96 DNA and Viral NA Small Volume Kit automated on the Magna Pure 96 (Roche, Switzerland) (SSI and L&F). The extracted RNA was used as template for a reverse transcriptase quantitative PCR (RT-qPCR) assay targeting the matrix gene of IAV (19,20) at SSI and DTU, and the ViroReal Kit Swine Influenza A (SIV) (Ingenetix, Austria) was used at L&F. All IAV positive submissions were initially screened for the presence of an H1pdm09 HA gene and thereafter the HA-NA lineage was determined for all submissions by multiplex RT-qPCR at SSI as previously described (16).

### Virus isolation in cells

Viruses were isolated by inoculation of MDCK-SIAT1 cells with clinical material using procedures described in the Manual for the laboratory diagnosis and virological surveillance of influenza, WHO Global Influenza Surveillance Network (21), with minor modifications (22).

### Whole genome sequencing

Each year a selection of samples (one sample per submission) with a Ct value ≤ 30 (25-89 samples per year) were selected for whole genome sequencing. The RNA was either obtained from the primary sample (nasal swab, oral fluid or lung tissue) or from viral isolates and used as template for a conventional PCR with the universal IAV primers “MBTuni-12R” and “MBTuni-13” and the resulting PCR products were sequenced using the Illumina MiSeq platform (23,24).

### Consensus generation and genotype assignment

Fastq files obtained from the Illumina Miseq platform were processed using “the SSI Influenza surveillance pipeline”. As part of the first step, the paired-end reads were initially filtered using fastp v0.20.1 (Chen et al., 2018) with the following parameters: ‘--trim_poly_g’, ‘--poly_g_min_len 7’, ‘--cut_tail’, ‘--cut_front --cut_window_size 6’, ‘--low_complexity_filter’ and ‘--complexity_threshold 50’. Subsequently, the filtered fastq files were used as input in the software KMA v1.3.27 (Clausen et al., 2018) to perform an iterative assembly of the influenza segments. The result of the assembly was compared against the NCBI influenza sequences database and, for each sample, the assembly with the highest coverage, depth and identity was selected.

The consensus sequences were then gathered and the command line version of Nucleotide-Nucleotide BLAST v2.12.0+ (Camacho et al., 2009) was used to identify the closest reference from a set of sequences representing each segment of all lineages present in the European swine population: H1N1pdm09 (1A.3.3.2), H1N2uk (1.B), Eurasian avian like H1N1 (1.C), Eurasian swine H3N2 and human seasonal H3N2. The HA sequenced were further classified into different lineages using the Swine influenza H1/H3 (global classification) tool at https://www.bv-brc.org/app/SubspeciesClassification.

### Genetic analysis

Nucleotide and translated amino acid sequences were aligned using the MUSCLE algorithm in CLC main workbench version 22 (25). The receptor binding sites (RBS) and antigenic sites (Sa, Sb, Ca1, Ca2 and Cb) (26) of the H1 were manually annotated to the amino acid sequences of the H1pdm09Nx viruses for comparison between clusters. Similarly, the antigenic sites of the NA protein (27) were annotated to the HxN1av viruses for comparison between clusters.

Bayesian phylogenetic trees were generated for the HA gene sequences of H1pdm09Nx Danish swIAV surveillance and H1pdm09Nx reference sequences using MrBayes version 3.2.7a with the setting nst = mixed rates = invgamma (28,29).

HA genes from H1pdm09Nx viruses sequenced through the Danish swIAV surveillance in 2013-2018 and previously analyzed (Ryt-Hansen et al., 2021) were added to the 2019-2022 H1pdm09NX dataset to infer phylogenetic trees using a strict molecular clock model in BEAuti and BEAST2 version 2.5.2 (30) as previously described (Ryt-Hansen et al., 2021), following an assessment of the temporal signal using TempEst (Rambaut et al., 2016). The strict molecular clock trees were used to estimate the evolutionary rate and the time of the novel H1pdm09N1av reassortant emergence. The program Tracer version 1.7.1 was used for visualizing the convergence and probabilities (31).

The PAML package was used to evaluate the presence of positive selection acting on evolution and amino acid positions prone to positive selection (32). In addition, the transient selection occurring on codons was also investigated using the HyPhy MEME package and web application Datamonkey with a p-value threshold of 0.05 (33,34). PAML was also used to map mutations onto branches of the phylogenetic trees, and ancestral reconstruction was run using the setting Rateancestor = 1. Phylogenetic trees were visualized in FigTree version 1.4.4 (35).

### Viral molecular markers related to an increase in the zoonotic potential of swIAV

Sequences and phylogenetic trees were searched for the presence of viral molecular markers indicative of enhanced zoonotic potential. Absence of the PB2 E627K mutation, implicated in the replication of avian IAVs in humans (36), in H1N1pdm09 is compensated by PB2 271A, 590S and 591R (37). Mutations in PA proteins that can enhance the pathogenicity and transmission of IAV in ferrets are V100I, N321K, I330V and A639T (38). MxA antiviral resistance is enhanced by NP protein mutations 48Q, 53D 98K, 99K, 100I/V and 313V (4). NA proteins can acquire resistance against neuraminidase inhibitors with mutations H275Y and N295S in N1 subtypes and R292K and E119G/D/A/V in N2 subtypes. Truncation of the NS1 protein from H1N1av origin was observed in A/Denmark/1/2021(H1N1v) (39) and similar NS1 truncation has been related to increased fitness of avian viruses in human cells (40).

### Antigenic characterization as determined by hemagglutination inhibition (HI)

Eight H1pdm09N1av viruses were tested against ferret antisera raised against five human seasonal H1N1pdm09 vaccine/circulating strains and one swine hyperimmune serum raised against the swine H1N1pdm09 vaccine “Respiporc FLUpan H1N1” (Ceva Animal Health). Initially the HA titer of each viral isolate was determined and four HA units of the viral isolate along 0.65 % guinea pig RBC was used for the HI-test as previously described (41). Additionally, the sera were first treated using a receptor destroying enzyme (RDE).

### Transmission study in ferrets

The ferret studies were performed in Biosafety Level 2 facilities St. Jude Children’s Research Hospital, Memphis, USA. H1pdm09N1av viruses representing the two different genotypes circulating in Denmark were selected: A/swine/Denmark/19922-5/2021, an H1pdm09N1av virus showing high nucleotide identities (99-99.9 %) to a Danish zoonotic case (18) and having a NS gene of Eurasian avian-like origin (H1pdm09N1av - Genotype 2) and A/swine/Denmark/15063-1/2020, an H1pdm09N1av reassortant having an NS gene of H1N1pdm09 origin (H1pdm09N1av - Genotype 1). The nucleotide identity between the two inoculum strains and the Danish H1pdm09N1av zoonotic case “A/Denmark/36/2021” are listed in Supplementary File 1.

The swIAV isolates were isolated in MDCK-SIAT cells and diluted in sterile PBS to give a concentration of the inoculum of 10^5^ 50% tissue culture infectious dose (TCID_50)_/mL as determined by the method of Reed and Muench (42).

The two ferret studies were performed in separate cubicles. For each viral strain, three donor ferrets were inoculated on day 0 with 1mL of virus (10^5^ TCID_50_) and were housed in separate cages. At day 1 (24 hours later), direct contacts were co-housed with the donors, and airborne contacts were housed in the same cage but separated physically from the donors and the direct contacts. This study design resulted in nine ferrets per viral strain including one donor ferret, one direct transmission ferret and one aerosol contact per cage. The ferrets were weighed and had their temperature measured by use of a subcutaneously injected microchip (Bio Medic Data Systems, Waterford Wisconsin, USA) daily from days 0-10 and at day 13. Pyrexia was defined as a body temperature > 40°C. In addition, a clinical score was given at the same time points in regard to their level of activity, body condition, dehydration, neurological distress and respiratory distress. At day 2, 5, 7 and 9 nasal washes were performed following administration of 0.4 mL of ketamine I.M as previously described (43). At day 14, all ferrets were euthanized and tissues from the nasal turbinates, the left (LU 1) and right (LU4) cranial lung lobes and the caudal right lung lobe (LU9) were collected. In addition, a serum sample was obtained from each ferret. The nasal washes and tissues were stored at −80 until viral titration in MDCK cells. Viral titers were determined by inoculating 0.1 mL of 10-fold dilutions of virus onto MDCK monolayers in 96-well tissue culture plates and then calculating TCID_50_ titers by the method of Reed and Muench (42). The lower limit of virus detection was 1.0 log10 TCID50/mL or 1.0 log10 EID50/mL. The serum samples were stored at −20 until being analyzed for the presence of homologous HA antibodies to the inoculum strains using HI-test as previously described.

### Level of preexisting immunity in the public

To test the population immunity towards the two H1pdm09N1av strains (A/swine/Denmark/19922-5/2021 and A/swine/Denmark/15063-1/2020) that were also evaluated for transmission in ferrets, 119 human sera representing different age groups were purchased from BioIVT, Westbury, NY. The sera was divided into six groups according to the age of the person from which they were obtained. Each age-group consisted of sera from 19-21 individuals of different sex and race (Supplementary File 2). The sera was tested in the HI-test (as described previously) for cross-reactive antibodies towards the two H1pdm09N1av strains described above. In addition, four control sera from WHO (WHO Reference Serum H1a G.57 a-A/PR/8/34, WHO Reference Serum H1b G.40, 41, 42, a-A/F.M./1/47, WHO Reference Serum H1c G.7 a-A/Swine//IA/15/30 and WHO Reference Serum H1d G.704 (2010) a-A/Duck/Alberta/35/76) was included. A titer ≥ 40 was regarded positive.

## Results

### Screening for swIAV and HA genes of H1N1pdm09 origin

The Danish swIAV surveillance program screened a total of 8791 samples from 2616 submissions received from 2019 through 2022 (586, 723, 857 and 487 submissions per consecutive year). Submissions were collected from 391-753 swine herds each year, representing 15-26 % of all swine herds in Denmark (44).

At least one swIAV positive sample was detected in 53-55 % of submissions received in 2019-2021 and in 68 % of submissions in 2022 (Figure 1A). The proportion of swIAV positive submissions with HA genes of H1N1pdm09 origin similarly increased from 20 to 37 % (2019–2022) with a peak of 42 % observed in 2021.

**Figure 1.**
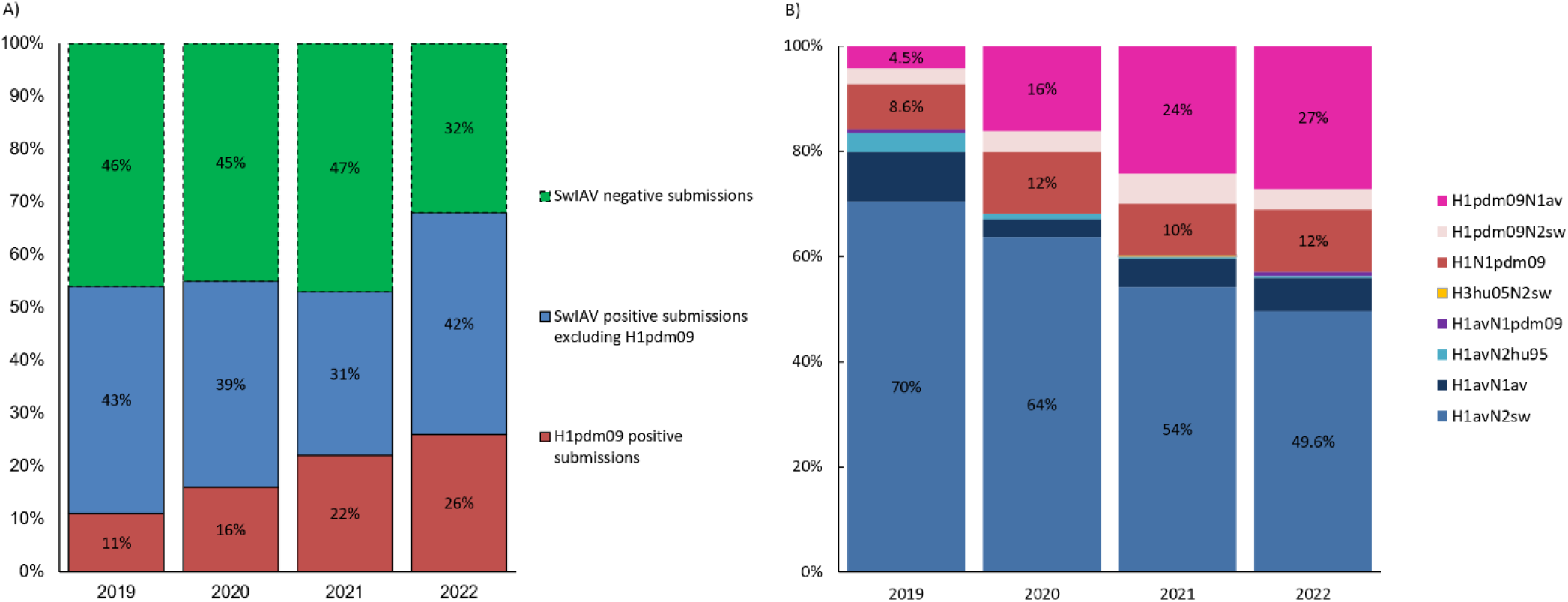
Proportion of IAV positive/negative and H1pdm09 positive submissions (A) and the proportion of the different HA and NA lineage combinations (subtypes) (B) registered from 2019-2022. 1B) The percentage of the H1pdm09N1av, H1N1pdm09 and H1avN2sw are highlighted on the columns to illustrate the relation between the increase of H1pdm09N1av with the decrease of H1avN2sw, while the proportion of H1N1pdm09 remains stable.

### Lineage determination and prevalence

The HA-NA lineages were determined for 44-64 % of the swIAV positive submissions per year by RT-qPCR, showing a fluctuating presence of eight swIAV HA and NA combinations (Figure 1B). Most strikingly, the annual distribution of the lineages revealed a major increase in the proportion of H1pdm09N1av viruses from 4.5% (2019) to 27 % (2022), which coincided with a decrease of H1avN2sw from 70% (2019) to 49.6 % (2022), while the proportion of H1N1pdm09 remained stable at around 10 %. Less common swIAVs were also observed including nine samples of “H1avN2hu95” and a single detection of “H3hu05N2sw” in 2021 bearing similarity to previous detections from 2013-2015 (45). For the H1 sequences of the H1av lineage, the majority of the sequences belonged to the 1C.2.4 clade (86/115), some to the 1C.2 and 1C.5 clades (10/115 and 11/115, respectively) and few to the 1C.2.1 and 1C.2.2 clades (3/115 and 5/115 respectively). All of the H1 sequences of the H1pdm09 lineage belonged to the 1A.3.3.2 clade, whereas the single H3 sequence belonged to the 2000.3 clade.

### Genotypic composition

A selection of the subtyped samples (n = 218) was subsequently sequenced to determine the lineage of the “internal gene cassette” (i.e. non-HA and NA gene segments) and genotypic composition (Figure 2). The sequences are available at NCBI GenBank with accession numbers: PQ115369 - PQ117065 (a detailed sequence list is provided in Supplementary File 3). For H1avN2sw viruses, the dominant genotype (Genotype 1: 66 % of H1avN2sw sequenced samples) had the full internal gene cassette of H1N1pdm09 origin (PPPPPP). The second most prevalent genotype (Genotype 2: 9 % of H1avN2sw sequenced samples) had the internal gene cassette of H1N1pdm09 origin with an NS gene segment of H1N1av origin (PPPPPA).

**Figure 2.**
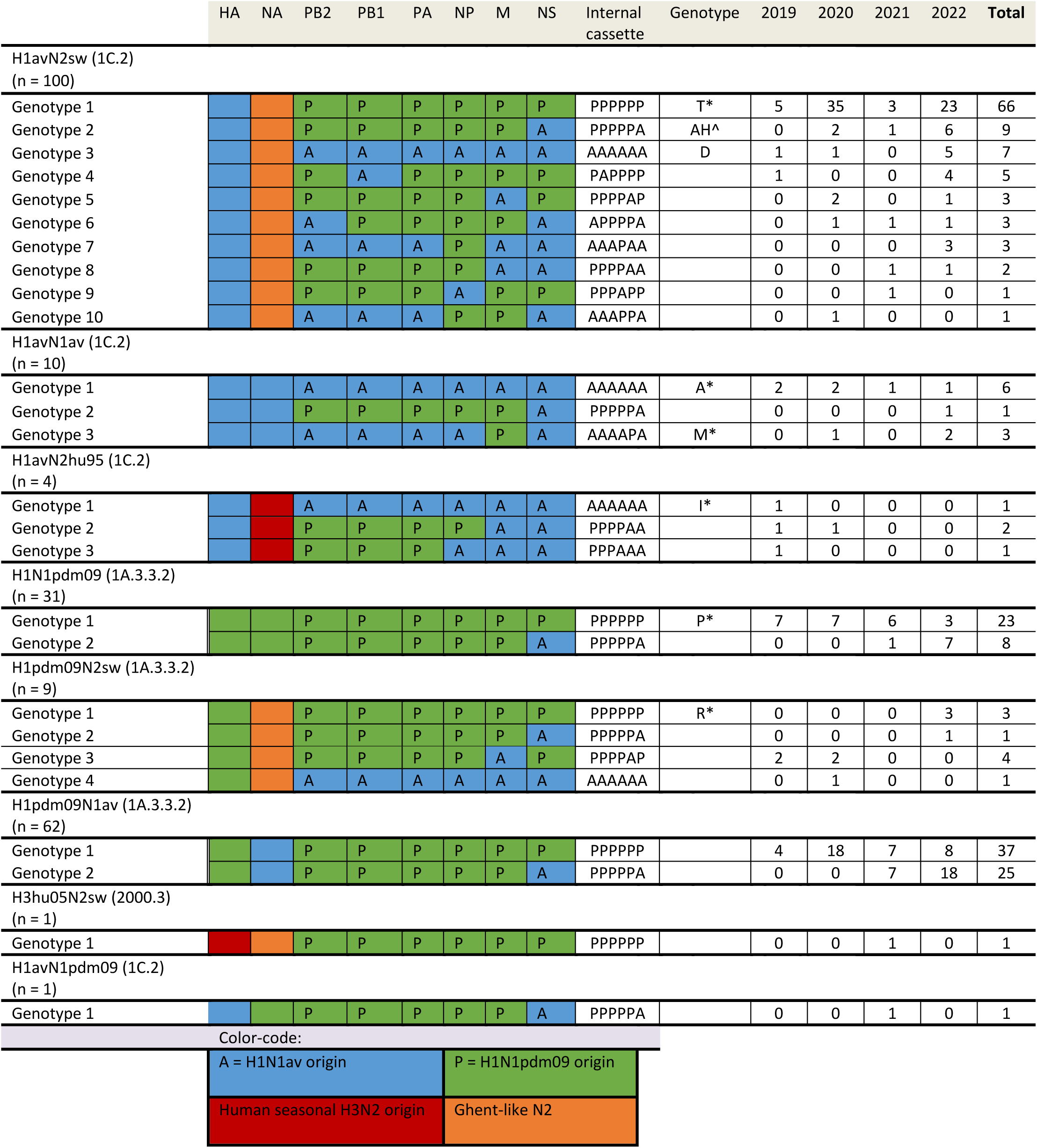
Genotypes of the Danish swIAV detected between 2019-2022. The origin of each gene segment “HA, NA, NP, NS, PA, PB1 and PB2” is indicated by the color and a letter for the internal genes, where A = Eurasian avian-like H1N1 (H1N1av) origin and P = H1N1pdm09 origin. The column “genotype” refers to the genotype according to the terminology defined by two previous studies by Wastson et al., 2015 and Henritzi et al., 2020. The number of each genotype is presented for the years 2019-2022 and as a total in the last column.

For H1avN1av viruses, the dominant genotype (Genotype 1: 60 % of H1avN1av sequenced samples) had the full internal gene cassette of H1N1av origin (AAAAAA). Less prevalent genotypes had the internal gene cassette of H1N1av origin with the M gene segment of H1N1pdm09 origin (AAAAPA) (Genotype 2: 30 % of H1avN1av sequenced samples) or the internal gene cassette of H1N1pdm09 origin with an NS gene segment of H1N1av origin (PPPPPA) (Genotype 3: 10 % of H1avN1av sequenced samples).

For H1N1pdm09 viruses, two genotypes were present; one (Genotype 1: 74 % of H1N1pdm09 sequenced samples) with the full H1N1pdm09 internal gene cassette (PPPPPP) and the other (Genotype 2: 26 % of H1N1pdm09 sequenced samples) with the internal gene cassette of H1N1pdm09 origin with an NS gene segment of H1N1av origin (PPPPPA). Interestingly, Genotype 2 became the dominating genotype by 2022 accounting for 70 % of H1N1pdm09 viruses sequenced that year.

For H1pdm09N1av viruses, two genotypes were also present; one (Genotype 1: 60 % of H1pdm09N1av sequenced samples) with the full H1N1pdm09 internal gene cassette (PPPPPP) and the other (Genotype 2: 40 % of the H1pdm09N1av sequenced samples) with the internal gene cassette of H1N1pdm09 origin with an NS gene segment of H1N1av origin (PPPPPA). Genotype 2 was first detected in 2021 and became the dominant genotype constituting 69 % of H1pdm09N1av viruses sequenced in 2022.

Overall, the internal cassette of PPPPPA became increasingly prominent over the four years going from undetected in 2019 to 38 % of genotyped samples by 2022; this combination was found in six of the eight HA/NA combinations detected.

### Phylogenetic analysis of the HA gene from H1pdm09Nx viruses

The Bayesian phylogenetic tree inferred from the HA genes of Danish swIAV H1pdm09 viruses isolated from 2019-2022 and reference sequences revealed three main clusters (Figure 3). One cluster (“swine adapted H1pdm09Nx viruses”) contained H1N1pdm09 and H1pdm09N2sw viruses, another (“H1pdm09N1av reassortants”) contained H1pdm09N1av viruses and a third (“human seasonal-like H1pdmNx viruses”) that contained swIAV H1N1pdm09 viruses that clustered more closely with human seasonal H1N1pdm09 viruses. The two major swine-adapted clusters, “swine adapted H1pdm09Nx viruses” and “H1pdm09N1av reassortants”, further split into sub-clusters that segregated with the origin of the NS gene segment.

**Figure 3.**
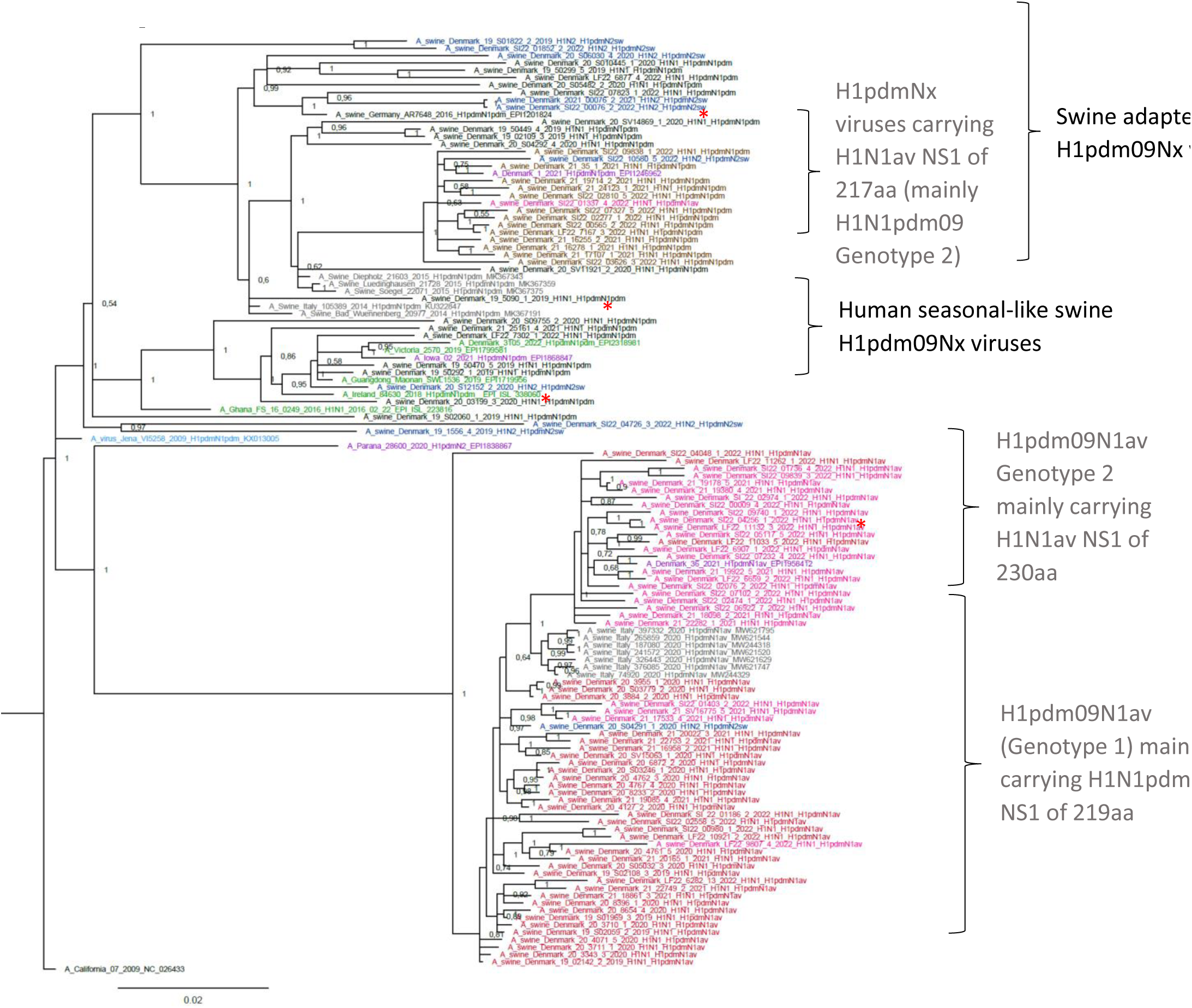
Bayesian phylogenetic tree of Danish H1pdm09Nx sequences and H1pdm09Nx reference sequences. A_California_07_2009 was used as an outgroup. Node labels represent posterior probabilities. Red and pink taxon = Danish H1pdm09N1av reassortant; red with full internal gene cassette of H1N1pdm09 origin and pink similar but with an Eurasian avian-like NS gene segment. Blue = Danish H1pdm09N2sw viruses. Black and brown = Danish H1N1pdm09 viruses; black with full internal cassette of H1N1pdm09 origin and brown similar but with an Eurasian avian-like NS gene segment. Grey = other European swine H1pdm09Nx viruses clustering with Danish H1pdm09Nx viruses. Purple taxon and red star = recent human zoonotic cases. Green = human H1N1pdm09 seasonal strains from 2016-2022. Light blue = swine H1N1pdm09 vaccine strain of Respiporc FluPan H1N1 (Ceva Animal Health).

### Evolutionary rate of the H1pdm09Nx HA gene

The HA gene sequences from H1pdm09Nx viruses were extracted from the 2019-2022 sequencing dataset and added to a previously studied dataset containing HA gene sequences from a selection of H1pdm09 viruses sequenced in 2013-2018 (16). The 2013-2022 sequence dataset gave a strong temporal signal with a correlation coefficient of 0.91, supporting phylogenetic analysis using a strict molecular clock model. Interestingly, the phylogenetic trees showed that the H1pdm09N1av cluster diversified after 2018, the same year in which the first H1pdm09N1av reassortant was detected thereby segregating with the NA type (Figure 4). The divergence of the two HA sub-clusters, which coincided with NS reassortment, was estimated to have occurred in the beginning of 2020.

**Figure 4.**
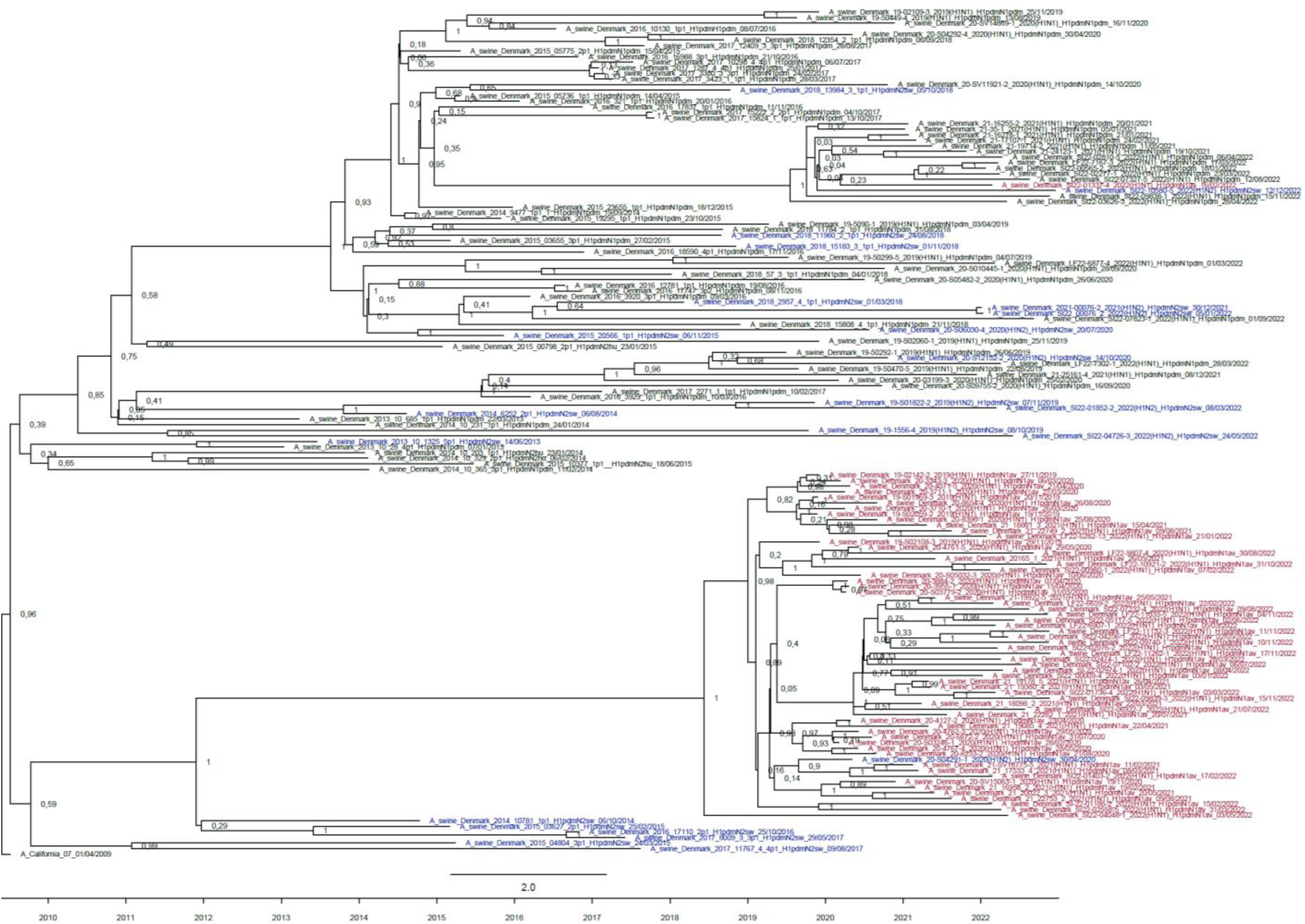
Strict molecular clock tree of the Danish H1pdm09 sequences derived from H1pdm09Nx viruses collected during the swIAV surveilance from 2013-2022. Node labels represent posterior probabilities. Black taxons indicate H1N1pdm09 viruses, blue taxons indicate H1pdm09N2sw viruses and red taxons indicate H1pdm09N1av viruses. The timeline from 2009-2022 is indicated below.

The estimated evolutionary rate of the 2013-2022 H1pdm09Nx HA gene was estimated to be 4.9 x 10^-3^ (SEM: 1.06 x 10^-5^) nucleotide substitutions per site per year. To measure if the rapid diversification of H1pdm09N1av affected the evolution rate, molecular clock analysis was repeated specifically on the HA genes of H1pdm09N1av reassortant viruses. The evolutionary rate of the H1pdm09N1av HA gene was significantly (p-value < 0.001) elevated at 5.35 x 10^-3^ (SEM: 9.23 x 10^-6^) nucleotide substitutions per site per year.

### Positive selection in H1pdm09Nx proteins

Positive selection was present in fourteen different sites in the HA of the H1pdm09Nx viruses (Supplementary File 4). In total, 13/14 of these sites were located in the RBS and/or in specific antigenic sites of the HA protein. An additional six sites were identified as being under transient positive selection and included 113, 158 (in the RBS), 187 (in the RBS and Ca1), 210 (in the RBS and Sb), 212 (in the RBS and Sb) and 405 (numbering from the first Methionine).

### Mutations unique to the H1pdm09N1av reassortant viruses

The viruses of the H1pdm09N1av phylogenetic cluster showed 31 unique HA mutations compared to the remaining H1pdmNx viruses in the tree (Supplementary File 5). The majority of these mutations (64 %) were located in the RBS and/or in specific antigenic sites. Similarly, the NA protein of the H1pdm09N1av viruses showed 14 unique mutations with four being located in antigenic sites (Supplementary File 6). The NA proteins of the H1pdm09N1av viruses showed an overall amino acid identity of 97-100 % to each other, whereas they shared 86-89% identity to the circulating H1N1av stains

### Viral molecular markers of zoonotic potential

Alignments of all 2019-2022 Danish swIAVs were examined for genetic markers known or suspected to increase viral fitness of swIAVs in humans.

The PB2 271A, 590S and 591R combination was prominent in the H1N1pdm09 viruses and was observed in 59 % of viruses. Substitutions at PB2 590 were noted in H1pdm09Nx viruses, with S590N inherited by 15 % of PB2 genes, including A/Denmark/1/2021(H1N1v), and S590G adopted in 11 % of PB2 genes, including A/Denmark/36/2021(H1N1v).

The PA V100I, N321K, I330V and A639T substitutions were present in ten viruses that included H1avN2sw (n = 7), H1N1pdm09 (n = 2) and H1avN1pdm09 (n = 1). Interestingly, the two H1N1pdm09 viruses (A/swine/Denmark/LF22-7302-1/2022(H1N1) and A/swine/Denmark/19-50470-5/2019(H1N1)) were likely reverse zoonotic IAVs and were most closely related to human seasonal H1N1pdm09 strains (Figure 3 – Human seasonal-like swine H1pdm09Nx viruses), while A/Denmark/1/2021(H1N1v) and A/Denmark/36/2021(H1N1v) both expressed PA 100V, 321K, 220V and 639T.

The NP mutations (48Q and 98K) that have been linked to MxA resistance were found in all H1N1av origin NP genes, whereas the 99K mutation was present in 20/21 H1N1av NPs. The NP mutations 53D, 100I/V and 313V, also linked to MxA resistance, were observed in 15 % (30/197) of the H1N1pdm09 origin NP proteins. Two of the three mutations (100I/V and 313V) were present in 68.5 % (135/197) of the H1N1pdm09 origin NP proteins whereas the remaining NP proteins only carried one of the two mutations (100I/V or 313V). In conclusion, all H1N1av and H1N1pdm09 NPs carried at least one of the MxA-resistance mutations.

NS1 proteins of varying lengths were detected in the Danish swIAVs. For the H1pdm09Nx viruses, NS lengths according to genotype are listed in Supplementary File 7 and are also indicated by the taxon color in Figure 3. The longest NS1 proteins, at 230 amino acids, were detected in most NS genes of H1N1av origin and is expressed by the viruses carrying a complete H1N1av internal gene cassette (AAAAAA) (for example H1avN2sw, H1N1av and H1avN2hu95 viruses). The cluster of H1pdmN1av genotype-2 viruses containing a PPPPPA internal gene cassette (including the second Danish zoonotic case (A/Denmark/36/2021(H1N1v)), all carry this version of the NS1 protein. In addition, other viruses with a similar internal gene cassette (PPPPPA) also carry a complete 230 amino acid NS1 protein with the exception of six viruses (4: H1avN2sw, 1: H1pdm09N2sw, 1: H1pdm09N1av) that had the 217 C-terminally truncated version of the H1N1av NS1 protein. Interestingly, the same 217 C-terminally truncated version of NS1 was expressed by the genotype 2 H1N1pdm09 viruses, which includes the first Danish zoonotic case virus A/Denmark/1/2021(H1N1v). C-terminal truncation of NS1 to 219 amino acids is typical of H1N1pdm09 viruses and was present in most of the viruses carrying a complete H1N1pdm09 gene cassette including H1avN2sw, H3hu05N2sw, H1pdm09N1av genotype 1 and H1N1pdm09 genotype 1 viruses. Other, less frequent, versions of the NS1 proteins, included those in three H1avN2sw viruses with a PPPPPA internal gene cassette with a 220 C-terminally truncated NS1 and one H1pdm09N2sw virus (A/swine/Denmark/6030-4/2020) with a PPPPPP internal gene cassette with a NS1 230 amino acids long. Thus, several versions of the NS1 gene are present in Danish swIAVs

### Antigenic characterization of H1pdm09N1av reassortant viruses and pre-exiting immunity in the public

A selection of eight H1pdm09N1av viruses were assessed for reactivity with post-infection ferret antisera raised against human seasonal H1N1pdm09 vaccine viruses in HI assay. No cross reaction was observed to the ferret antisera representing the human seasonal H1N1pdm09, with the exception of five strains reacting slightly with ferret sera raised against A/California/07/09 (maximum titer of 40) (Table 1). All eight viruses H1pdm09N1av showed some level of cross reaction to the hyperimmune sera raised against the swine influenza vaccine Respiporc FLUpan H1N1 with titers ranging from 80-320. However, the cross-reaction between humans strain similar to that used in the H1N1pdm09 swine vaccine, showed much higher cross-reaction with titers reaching 1280 for example to A/California/07/09 and A/Hamburg/1580/2009.

**Table 1.** HI titers of H1pdm09N1av isolates. Red = positive titers. Green = negative (<20). Grey = cross reference and sera. The H1pdm09N1av isolates were tested against ferret antisera towards human seasonal H1N1pdm09 vaccine strains (A/Sydney/5/2021 F34/22 egg RDE INF172 01-05-24 ISHY, A/Victoria/2570/2019 F26/20 egg 12-11-2024 MIWP, A/California/07/09 (C4/16/09) RDE: 06-11-2024 SAMD, A/Michigan/45/15 (F40/19) RDE: 11-01-2024 ISHY, A/Brisbane/02/2018 F10/19 egg RDE:11-01-24 ISHY) from WHO and swine hyperimmune raised towards the H1N1pdm09 swine vaccine Respiporc FLUpan (Ceva Animal Health) (Pan H1N1 animal 5976 TV 1875/19 RDE: 22-05-24 DALP). A/Victoria/2570/19 was used as a positive control.

|  | A/Sydney/5/2021 | A/Victoria/2570/2019 | A/California/07/09 | A/Michigan/45/15 | A/Brisbane/02/2018 | Respiporc FLUpan |
| --- | --- | --- | --- | --- | --- | --- |
| **Isolate:** |  |  |  |  |  |  |
| A/Sw/DK/15063-1/2020 |  |  | 40 |  |  | 160 |
| A/Sw/DK/16775-5/2021 |  |  | 20/40 |  |  | 80 |
| A/Sw/DK/17023-1/2021 |  |  |  |  |  | 160 |
| A/Sw/DK/19537-2/2021 |  |  |  |  |  | 160/320 |
| A/Sw/DK/19922-1/2021 |  |  | 40 |  |  | 160 |
| A/Sw/DK/01337-4/2022 |  |  | 40 |  |  | 160 |
| A/Sw/DK/01403-2/2022 |  |  |  |  |  | 80/160 |
| A/Sw/DK/04256-1/2022 |  |  | 40 |  |  | 80 |
| A/Victoria/2570/2019 | 640 | 1280 | 20 | 40 | 20 | 160/320 |
| **Reference strains:** |  |  |  |  |  |  |
| A/Sydney/5/2021 | 320 | 640 | 20 | 20 | 20/40 | 80 |
| A/Victoria/2570/2019 | 640 | 1280 | 20 | 40 | 20/40 | 320 |
| A/California/07/09 | 40 | 80 | 160 | 160 | 80 | 1280 |
| A/Michigan/45/2015 | 20 | 40 | 160 | 320 | 320 | 1280 |
| A/Brisbane/02/2018 | 20 | 40 | 320 | 320 | 640 | 1280 |
| A/Hamburg/1580/2009 | 20 | 20 | 320 | 320 | 320 | 1280 |

Human sera from 119 persons was obtained to test the level of preexisting immunity towards the two H1pdm09N1av strains (A/swine/Denmark/19922-5/2021 and A/swine/Denmark/15063-1/2020) representing the two genotypes of H1pdm09N1av viruses and the same viruses selected for the ferret transmission study. Of the serum from 119 individuals of different ages, 21 (17.6 %) had HI titers ≥40 against the A/swine/Denmark/19922-5/2021 strain, while only four (3.3 %) had HI titers ≥40 against the A/swine/Denmark/15063-1/2020 strain. As presented in Figure 6 it is observed that the youngest (0-18 years) and oldest persons (66+) has the highest proportion of HI positive sera against A/swine/Denmark/19922-5/2021, while none of sera representing the younger population (0-25 years) have any cross-reactive antibodies towards A/swine/Denmark/15063-1/2020. The level of antibodies in the individual HI positive sera against A/swine/Denmark/19922-5/2021 ranged between 40-1280. On the contrary the level of antibodies in the HI positive sera against A/swine/Denmark/15063-1/2020 was 80 at the highest.

### Successful aerosol transmission of the H1pdm09N1av genotype 1 in ferrets

A/swine/Denmark/19922-5/2021, an H1pdm09N1av - genotype 2 virus, and A/swine/Denmark/15063-1/2020, an H1pdm09N1av - genotype 1 virus, were inoculated intranasally into ferrets. Nasal washes of all inoculated donor ferrets were IAV positive 2 days post inoculation (DPI) and viral titers either remained stable or increased slightly by 5 DPI (Figure 5). All donor ferrets inoculated with A/swine/Denmark/19922-5/2021 had cleared virus by 7 DPI, whereas 2/3 ferrets inoculated with A/swine/Denmark/15063-1/2020 were still positive at 7 DPI. All direct contact ferrets, introduced into the same cage as the donor ferrets on 1 DPI, were also IAV positive at 2 DPI and remained IAV positive through 7 DPI (four of the six direct contact animals had cleared virus by 9 DPI). Airborne contacts were introduced to an adjacent cage on 1 DPI. Airborne contacts of A/swine/Denmark/15063-1/2020 infected donors displayed no clear evidence of infection with only 1/3 ferrets (ferret #4102) testing weakly positive at 2 DPI (Figure 5A). However, the airborne contacts of A/swine/Denmark/19922-5/2021 donors showed clear evidence of transmission with 2/3 ferrets positive at 7 DPI and 3/3 ferrets positive at 9 DPI (Figure 5B).

**Figure 5.**
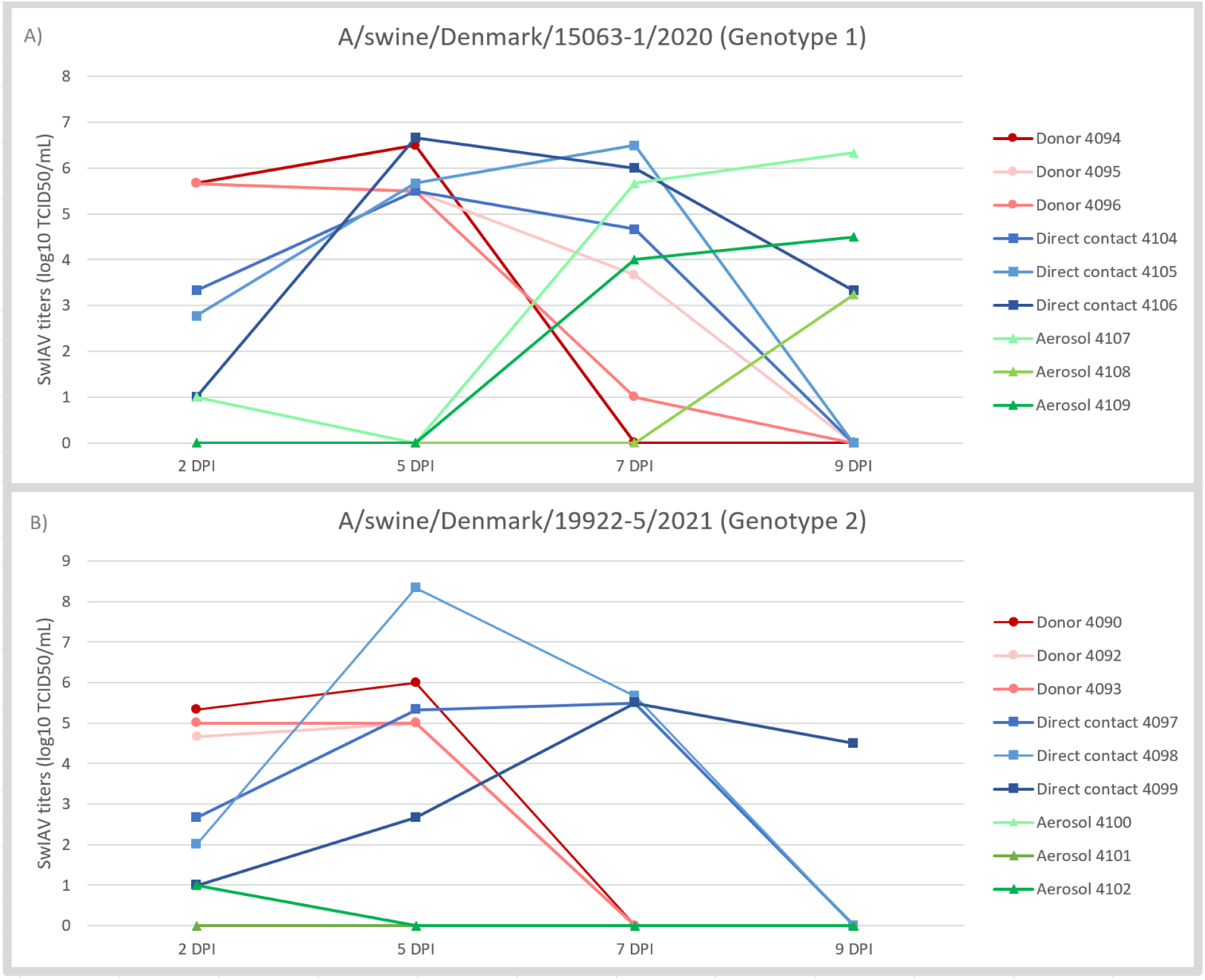
Viral titers of the nasal washes of the donors, direct- and aerosol contacts ferrets innoculated with the two different H1pdm09N1av strains; Genotype 1 (A) and Genotype 2 (B). It should be noted that “Donor 4092” have a similar decrease as “Donor 4093” on DPI 7. The exact titers of each ferret are included in Supplementary File 1.

**Figure 6.**
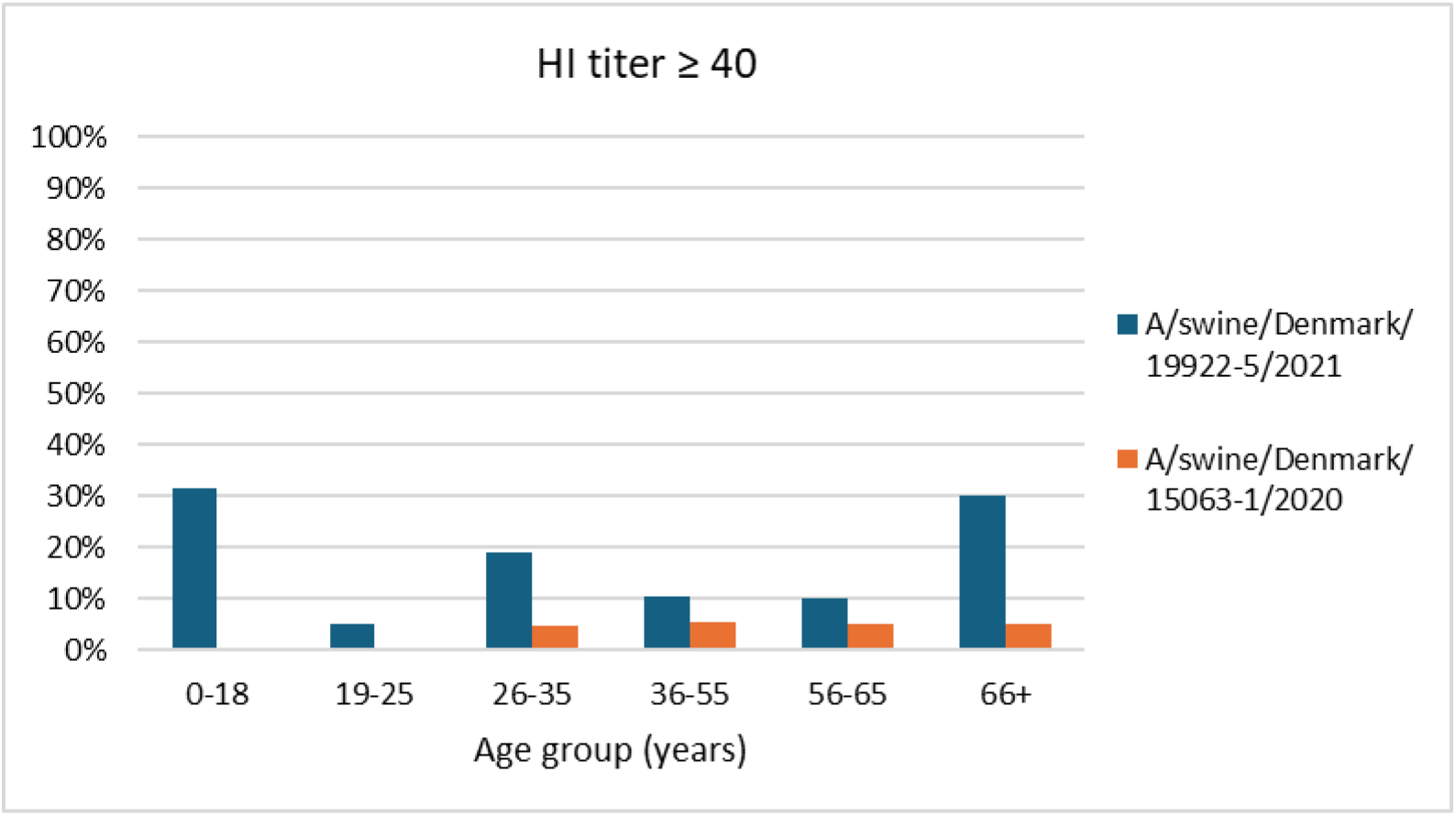
Level of pre-existing immunity against H1pdm09N1av genotype 1 and 2 in the general public. The figure represents the proportion of HI positive sera (HI titer ≥40) of the 119 sera from six different age-groups of the general public against the A/swine/Denmark/19922-5/2021 and A/swine/Denmark/15063-1/2020 strains, respectively.

Tissues obtained at 14 DPI were all virus negative by viral titration with the exception of one ferret in each inoculum group, which tested positive in the nasal turbinates (Supplementary File 8); the two positive animals were those that were virus positive in nasal washes at 9 DPI.

In the A/swine/Denmark/15063-1/2020 group, very few clinical signs were registered. One donor ferret and one direct contact ferrets showed pyrexia, in addition to two direct contact ferrets sneezing. More clinical signs were registered in the A/swine/Denmark/19922-5/2021 group, where lower activity levels, fever and diarrhea were observed in some of the donor and direct contact ferrets. In addition, one of the aerosol contacts showed sneezing at 8 DPI and decreased activity over the following three days as well as pyrexia at 8 and 9 DPI. By 11 DPI this ferret also had unkempt fur and dull eyes as well as nasal discharge and therefore was euthanized the same day (Supplementary File 9).

The average weight loss of the inoculated donor ferrets and direct contact ferrets from 0-13 and 0-7 DPI, respectively, was calculated and all ferrets lost weight during the study. No significant differences in weight loss were observed between the donor and direct contact ferrets of both inoculum groups. For the airborne contacts, weight loss was observed in all the ferrets of the A/swine/Denmark/19922-5/2021 group, consistent with virus isolation data. However, for the airborne contacts of the A/swine/Denmark/15063-1/2020 group, only one ferret exhibited a small weight loss without any sign of viral replication, while the other two ferrets gained weight (Supplementary File 9).

At 14 DPI all donor and direct contact ferrets had positive HI-titers (320–1280) towards the inoculum strains, whereas all aerosol contacts had HI-titers <10 (Supplementary File 10).

## Discussion

Passive IAV surveillance in swine in Denmark identified the rapid spread of a novel H1pdm09N1av reassortant virus from 2019-2022. This rapid spread suggests that these reassortant viruses have a selective advantage compared to other contemporary circulating swIAVs, a possibility further supported by the fact that detections of previously dominant Danish H1avN2sw viruses decreased for the first time in ten years (16,46). Although the exact background for the emergence of the H1pdm09N1av are unclear, their biologic properties offer some suggestions. For example, the genetic diversification of the HA of the H1pdm09N1av reassortants, with mutations occurring in antigenic sites, may provide an avenue for their escape of preexisting immunity generated through prior exposure to other H1pdm09Nx viruses. In support of this, we were able to show a significant reduction in HI-titers between selected H1pdm09N1av strains compared to strains more similar to those included in the swine vaccine. Of note, the decline in detections of the H1avN2sw viruses concomitant with the rise in H1pdm09N1av reassortant detections was not observed when the H1N1pdm09 virus started circulating in Denmark in 2010 (16). Factors influencing virus fitness include properties such as the HA/NA balance (47,48), HA pH stability (49), NA activity (50), receptor binding affinity (51) and the immune response of the host (52–54).

The role that the specific pairing of the H1pdm09 HA with the N1av NA had on virus emergence is as of yet undetermined. Preferential HA and NA pairing has been observed in swine influenza surveillance in the US and has driven the emergence and recession of viral lineages over time. Similarly to what we have observed, reassortment events in the US have been temporally linked to increased evolutionary rates and antigenic drift promoting the success of novel reassortant viruses (9,10). We found that the H1pdm09N1av reassortants form a unique HA phylogenetic cluster (with mutations mainly occurring in antigenic sites and the RBS) and had a higher evolutionary rate than the other H1pdm09Nx viruses. This could indicate that the receptor affinity has to change to match the NA.

Investigators studying swIAV evolution in the US have previously hypothesized that a specific NP gene played an important role in the success of novel US H3N2 reassortants (9,10). Enhanced transmissibility of the H3N2 viruses carrying this NP gene was later confirmed using reverse genetics (55). Altogether these studies support the idea that the combination of internal genes is important for the success of swIAVs. For the H1pdm09N1av genotype 2 viruses in Denmark, the acquisition of the H1N1av NS gene may have contributed to their increased detection relative to the genotype 1 viruses. H1pdm09N1av genotype 2 viruses containing the PPPPPA internal gene cassette appeared soon after (2021) the first detection of H1pdm09N1av genotype 1 viruses (2018) and now constitute the majority of detected H1pdm09N1av reassortants. The finding that the PPPPPA internal gene cassette has been integrated into six of the eight HA/NA combinations of viruses circulating in Denmark does imply a selective advantage for viruses possessing it. It should also be highlighted that both zoonotic cases with swIAV in Denmark had a PPPPPA internal gene cassette (17,18), however, with different lengths of the H1N1av NS1 protein. The NS1 protein is known to play a key role in antagonizing the host innate immune response and modulating host gene expression (56–58). Different NS1 proteins were identified in swIAV in this study, several of which included C-terminally truncated versions derived from H1N1av and H1N1pdm09 lineages. Truncation of the C-terminus of an avian H5N8 virus NS1 gene has previously been linked to blocking of apoptosis and interferon induction with a resulting increase in virulence in mice (40).

The genetic drift observed following the new HA and NA pairing of the H1pdm09N1av reassortants also led to a change in the antigenic properties of these viruses. The H1pdm09N1av reassortants showed minimal reactivity (titers ranging from <20-40) to ferret antisera raised against older and current human H1N1pdm09 vaccine strains. The H1pdm09N1av genotype 2 reassortant did cause a zoonotic infection in a recently vaccinated healthy person and resulted in severe disease consistent with this lack of reactivity (18). Unfortunately, it was not possible to propagate the specific zoonotic case virus to investigate it further. However, a highly similar virus was identified, isolated in cells and chosen for the ferret transmission study along with a H1pdm09N1av genotype 1 virus. The results revealed efficient transmission of both viruses among the donors and direct contacts. However, only the H1pdm09N1av genotype 1 virus transmitted to airborne contact animals, indicating that the pandemic potential of the zoonotic virus may be currently limited without further changes. The limited potential antibody protection in the human population of both strains tested for transmission in ferrets further emphasize that these strains could have an increased zoonotic potential. Notably, the H1pdm09N1av genotype 1 virus had extremely low cross-reaction to the sera in all age groups particularly the youngest people between 0-25 years of age. Interestingly, the H1pdm09N1av genotype 2 virus showed the highest proportion of cross-reactive sera in the youngest and oldest persons, which are normally at the highest risk developing severe disease due to influenza infections (59). However, a previous study of the 2009 H1N1 pandemic did find that older children and young adults were more likely to develop severe disease compared to the seasonal flu circulating at that time (60).

From a human risk assessment perspective, in addition to ferret transmissibility and antigenic characteristics, the presence of molecular markers associated with human adaptation is important. Ten Danish swIAV strains (both H1avNx and H1pdm09Nx) carrying all four residues in the PA protein (V100I, N321K, I330V and A639T) shown to increase the pathogenicity and transmission of Eurasian avian-like swine H1N1 viruses in ferrets were identified (38). These viruses should be prioritized for further phenotypic characterization to evaluate their zoonotic potential. Additionally, all the Danish swIAV characterized carried at least one- and the majority several-mutation/s in the NP protein linked to MxA resistance and elevated zoonotic potential. Mutations compensatory to the PB2 E627K mutation were also identified in 59 % of the swIAVs.

Together, our longitudinal data have shown that changes have occurred in the population of swIAV circulating in Danish swine with many viruses having traits associated with zoonotic threat. Continued selective pressures and assortment opportunities will only continue to diversify swIAV with more zoonotic infections likely. Continued monitoring of human diagnostic systems to ensure detection of variant H1 infections and risk assessments must remain a priority.

## Acknowledgement

The authors would like to acknowledge the Danish farmers and veterinarians that submitted samples for the Danish national passive swIAV surveillance program. In addition, the authors would like to acknowledge the funders of the project; the Danish veterinary and food administration [grant number: I2RG2: 2019-2022] and the Novo Nordisk Foundation [FluZooMark grant number NNF19OC0056326].

## Data availability statement

Supplementary files are available online.

## References

1. Rose N, Hervé S, Eveno E, Barbier N, Eono F, Dorenlor V, et al. Dynamics of influenza A virus infections in permanently infected pig farms: evidence of recurrent infections, circulation of several swine influenza viruses and reassortment events. Vet Res [Internet]. 2013 [cited 2017 Oct 18];44(1):72. Available from: http://www.veterinaryresearch.org/content/44/1/72

2. Ryt-Hansen P, Larsen I, Kristensen CS, Krog JS, Wacheck S, Larsen LE. Longitudinal field studies reveal early infection and persistence of influenza A virus in piglets despite the presence of maternally derived antibodies. Vet Res [Internet]. 2019 Dec 22 [cited 2019 May 27];50(1):36. Available from: https://veterinaryresearch.biomedcentral.com/articles/10.1186/s13567-019-0655-x

3. Simon-Grifé M, Martín-Valls GE, Vilar MJ, Busquets N, Mora-Salvatierra M, Bestebroer TM, et al. Swine influenza virus infection dynamics in two pig farms; results of a longitudinal assessment. Vet Res [Internet]. 2012 [cited 2017 Oct 18];43(1):24. Available from: http://www.veterinaryresearch.org/content/43/1/24

4. Henritzi D, Petric PP, Lewis NS, Graaf A, Pessia A, Starick E, et al. Surveillance of European Domestic Pig Populations Identifies an Emerging Reservoir of Potentially Zoonotic Swine Influenza A Viruses. Cell Host Microbe. 2020;

5. Markin A, Zanella GC, Arendsee ZW, Zhang J, Krueger KM, Gauger PC, et al. Reverse-zoonoses of 2009 H1N1 pandemic influenza A viruses and evolution in United States swine results in viruses with zoonotic potential. Su S, editor. PLOS Pathog [Internet]. 2023 Jul 27 [cited 2023 Sep 19];19(7):e1011476. Available from: https://journals.plos.org/plospathogens/article?id=10.1371/journal.ppat.1011476

6. Rajão DS, Walia RR, Campbell B, Gauger PC, Janas-Martindale A, Killian ML, et al. Reassortment between Swine H3N2 and 2009 Pandemic H1N1 in the United States Resulted in Influenza A Viruses with Diverse Genetic Constellations with Variable Virulence in Pigs. J Virol. 2017;

7. Chiapponi C, Prosperi A, Moreno A, Baioni L, Faccini S, Manfredi R, et al. Genetic variability among swine influenza viruses in Italy: Data analysis of the period 2017– 2020. Viruses [Internet]. 2022 Jan 1 [cited 2024 Jul 15];14(1):47. Available from: https://www.mdpi.com/1999-4915/14/1/47/htm

8. Chastagner A, Hervé S, Bonin E, Quéguiner S, Hirchaud E, Henritzi D, et al. Spatiotemporal Distribution and Evolution of the A/H1N1 2009 Pandemic Influenza Virus in Pigs in France from 2009 to 2017: Identification of a Potential Swine-Specific Lineage. García-Sastre A, editor. J Virol [Internet]. 2018 Sep 26 [cited 2019 Feb 5];92(24). Available from: http://jvi.asm.org/lookup/doi/10.1128/JVI.00988-18

9. Neveau MN, Zeller MA, Kaplan BS, Souza CK, Gauger PC, Vincent AL, et al. Genetic and Antigenic Characterization of an Expanding H3 Influenza A Virus Clade in U.S. Swine Visualized by Nextstrain. mSphere. 2022 Jun 29;

10. Zeller MA, Chang J, Vincent AL, Gauger PC, Anderson TK. Spatial and temporal coevolution of N2 neuraminidase and H1 and H3 hemagglutinin genes of influenza A virus in US swine. Virus Evol. 2021;7(2):1–13.

11. Ma J, Shen H, McDowell C, Liu Q, Duff M, Lee J, et al. Virus survival and fitness when multiple genotypes and subtypes of influenza A viruses exist and circulate in swine. Virology [Internet]. 2019 Jun 1 [cited 2023 Mar 7];532:30–8. Available from: https://pubmed.ncbi.nlm.nih.gov/31003122/

12. Taubenberger JK, Kash JC. Influenza Virus Evolution, Host Adaptation, and Pandemic Formation. Cell Host Microbe [Internet]. 2010 Jun [cited 2019 Sep 9];7(6):440–51. Available from: https://linkinghub.elsevier.com/retrieve/pii/S1931312810001721

13. Murcia PR, Hughes J, Battista P, Lloyd L, Baillie GJ, Ramirez-Gonzalez RH, et al. Evolution of an Eurasian Avian-like Influenza Virus in Naïve and Vaccinated Pigs. PLOS Pathog [Internet]. 2012 May [cited 2023 Mar 7];8(5):e1002730. Available from: https://journals.plos.org/plospathogens/article?id=10.1371/journal.ppat.1002730

14. Boni A, Vaccari G, Di Trani L, Zaccaria G, Alborali GL, Lelli D, et al. Genetic characterization and evolution of H1N1pdm09 after circulation in a swine farm. Biomed Res Int [Internet]. 2014 [cited 2023 Mar 7];2014. Available from: https://pubmed.ncbi.nlm.nih.gov/25025062/

15. Ryt-Hansen P, Pedersen AG, Larsen I, Kristensen CS, Krog JS, Wacheck S, et al. Substantial antigenic drift in the hemagglutinin protein of swine influenza a viruses. Viruses. 2020;

16. Ryt-Hansen P, Krog JS, Breum SØ, Hjulsager CK, Pedersen AG, Trebbien R, et al. Co-circulation of multiple influenza a reassortants in swine harboring genes from seasonal human and swine influenza viruses. Elife. 2021 Jul 1;10.

17. Nissen JN, George SJ, Hjulsager CK, Krog JS, Nielsen XC, Madsen T V., et al. Reassortant Influenza A(H1N1)pdm09 Virus in Elderly Woman, Denmark, January 2021. Emerg Infect Dis [Internet]. 2021 Dec 1 [cited 2022 Dec 6];27(12):3202–5. Available from: https://pubmed.ncbi.nlm.nih.gov/34808097/

18. Andersen KM, Vestergaard LS, Nissen JN, George SJ, Ryt-Hansen P, Hjulsager CK, et al. Severe Human Case of Zoonotic Infection with Swine-Origin Influenza A Virus, Denmark, 2021. Emerg Infect Dis [Internet]. 2022 Dec [cited 2022 Dec 6];28(12):2561–4. Available from: https://pubmed.ncbi.nlm.nih.gov/36418004/

19. Nagy A, Cerníková L, Kunteová K, Dirbáková Z, Thomas SS, Slomka MJ, et al. A universal RT-qPCR assay for “One Health” detection of influenza A viruses. PLoS One [Internet]. 2021 Jan 1 [cited 2022 Mar 22];16(1). Available from: https://pubmed.ncbi.nlm.nih.gov/33471840/

20. Nagy A, Vostinakova V, Pirchanova Z, Cernikova L, Dirbakova Z, Mojzis M, et al. Development and evaluation of a one-step real-time RT-PCR assay for universal detection of influenza A viruses from avian and mammal species. Arch Virol [Internet]. 2010 May 13 [cited 2017 Jun 12];155(5):665–73. Available from: http://link.springer.com/10.1007/s00705-010-0636-x

21. WHO. Manual for the laboratory diagnosis and virological surveillance of influenza [Internet]. 2011. Available from: https://iris.who.int/handle/10665/44518

22. Kristensen C, Laybourn HA, Crumpton JC, Martiny K, Webb A, Ryt-Hansen P, et al. Experimental infection of pigs and ferrets with “pre-pandemic,” human-adapted, and swine-adapted variants of the H1N1pdm09 influenza A virus reveals significant differences in viral dynamics and pathological manifestations. PLOS Pathog [Internet]. 2023 Dec 1 [cited 2024 Jul 18];19(12):e1011838. Available from: https://journals.plos.org/plospathogens/article?id=10.1371/journal.ppat.1011838

23. Liang Y, Nissen JN, Krog JS, Breum S, Trebbien R, Larsen LE, et al. Novel Clade 2.3.4.4b Highly Pathogenic Avian Influenza A H5N8 and H5N5 Viruses in Denmark, 2020. Viruses [Internet]. 2021 [cited 2024 Jul 8];13(5). Available from: https://pubmed.ncbi.nlm.nih.gov/34065033/

24. Kai Lee H. Simplified Large-Scale Sanger Genome Sequencing for Influenza A/H3N2 Virus. 2016 [cited 2019 Mar 19]; Available from: https://findit.dtu.dk/en/catalog/2341936806

25. C. Edgar R. MUSCLE: multiple sequence alignment with high accuracy and high throughput. 2013 [cited 2019 Jun 27]; Available from: https://findit.dtu.dk/en/catalog/2354275506

26. Manicassamy B, Medina RA, Hai R, Tsibane T, Stertz S, Nistal-Villán E, et al. Protection of Mice against Lethal Challenge with 2009 H1N1 Influenza A Virus by 1918-Like and Classical Swine H1N1 Based Vaccines. Fouchier RAM, editor. PLoS Pathog [Internet]. 2010 Jan 29 [cited 2019 Mar 7];6(1):e1000745. Available from: http://dx.plos.org/10.1371/journal.ppat.1000745

27. Lu X, Liu F, Zeng H, Sheu T, Achenbach JE, Veguilla V, et al. Evaluation of the antigenic relatedness and cross-protective immunity of the neuraminidase between human influenza A (H1N1) virus and highly pathogenic avian influenza A (H5N1) virus. Virology. 2014 Apr 1;454–455(1):169–75.

28. Ronquist F, Teslenko M, van der Mark P, Ayres DL, Darling A, Höhna S, et al. MrBayes 3.2: efficient Bayesian phylogenetic inference and model choice across a large model space. Syst Biol [Internet]. 2012 May [cited 2016 Sep 21];61(3):539–42. Available from: http://www.pubmedcentral.nih.gov/articlerender.fcgi?artid=3329765&tool=pmcentrez &rendertype=abstract

29. Ronquist F, Huelsenbeck JP. MrBayes 3: Bayesian phylogenetic inference under mixed models. Bioinformatics [Internet]. 2003 Aug 12 [cited 2019 Jul 3];19(12):1572– 4. Available from: https://academic.oup.com/bioinformatics/article-lookup/doi/10.1093/bioinformatics/btg180

30. Bouckaert R. BEAST 2: A software platform for Bayesian evolutionary analysis. 2016 [cited 2019 Jul 3]; Available from: https://findit.dtu.dk/en/catalog/2342383350

31. Rambaut A, Drummond AJ, Xie D, Baele G, Suchard MA. Posterior Summarization in Bayesian Phylogenetics Using Tracer 1.7. Susko E, editor. Syst Biol [Internet]. 2018 Sep 1 [cited 2019 Jul 10];67(5):901–4. Available from: https://academic.oup.com/sysbio/article/67/5/901/4989127

32. Yang Z. PAML 4: Phylogenetic Analysis by Maximum Likelihood. Mol Biol Evol [Internet]. 2007 Apr 18 [cited 2019 Jul 3];24(8):1586–91. Available from: https://academic.oup.com/mbe/article-lookup/doi/10.1093/molbev/msm088

33. Weaver S, Shank SD, Spielman SJ, Li M, Muse S V, Kosakovsky Pond SL, et al. Datamonkey 2.0: A Modern Web Application for Characterizing Selective and Other Evolutionary Processes.

34. Kosakovsky Pond SL, Poon AFY, Velazquez R, Weaver S, Hepler NL, Murrell B, et al. HyPhy 2.5—A Customizable Platform for Evolutionary Hypothesis Testing Using Phylogenies. Mol Biol Evol [Internet]. 2020 Jan 1 [cited 2024 Jul 15];37(1):295. Available from: /pmc/articles/PMC8204705/

35. Rambaut A. FigTree [Internet]. 2006 [cited 2019 Jun 6]. Available from: http://tree.bio.ed.ac.uk/software/figtree/

36. Zhu W, Li L, Yan Z, Gan T, Li L, Chen R, et al. Dual E627K and D701N mutations in the PB2 protein of A(H7N9) influenza virus increased its virulence in mammalian models. Sci Rep [Internet]. 2015 Sep 22 [cited 2022 Dec 6];5. Available from: https://pubmed.ncbi.nlm.nih.gov/26391278/

37. Liu Q, Qiao C, Marjuki H, Bawa B, Ma J, Guillossou S, et al. Combination of PB2 271A and SR polymorphism at positions 590/591 is critical for viral replication and virulence of swine influenza virus in cultured cells and in vivo. J Virol [Internet]. 2012 Jan 15 [cited 2024 Jul 8];86(2):1233–7. Available from: https://pubmed.ncbi.nlm.nih.gov/22072752/

38. Meng F, Yang H, Qu Z, Chen Y, Zhang Y, Zhang Y, et al. A Eurasian avian-like H1N1 swine influenza reassortant virus became pathogenic and highly transmissible due to mutations in its PA gene. Proc Natl Acad Sci U S A [Internet]. 2022 Aug 23 [cited 2022 Dec 6];119(34). Available from: https://pubmed.ncbi.nlm.nih.gov/35969783/

39. Nissen JN, George SJ, Hjulsager CK, Krog JS, Nielsen XC, Madsen T V., et al. Reassortant Influenza A(H1N1)pdm09 Virus in Elderly Woman, Denmark, January 2021. Emerg Infect Dis [Internet]. 2021 Dec 1 [cited 2022 May 19];27(12):3202–5. Available from: https://pubmed.ncbi.nlm.nih.gov/34808097/

40. Blaurock C, Blohm U, Luttermann C, Holzerland J, Scheibner D, Schäfer A, et al. The C-terminus of non-structural protein 1 (NS1) in H5N8 clade 2.3.4.4 avian influenza virus affects virus fitness in human cells and virulence in mice. Emerg Microbes Infect [Internet]. 2021 [cited 2022 Dec 6];10(1):1760–76. Available from: https://pubmed.ncbi.nlm.nih.gov/34420477/

41. Ryt-Hansen P, Pedersen AG, Larsen I, Krog JS, Kristensen CS, Larsen LE. Acute Influenza A virus outbreak in an enzootic infected sow herd: Impact on viral dynamics, genetic and antigenic variability and effect of maternally derived antibodies and vaccination. Tripp RA, editor. PLoS One [Internet]. 2019 Nov 14 [cited 2020 Jan 27];14(11):e0224854. Available from: https://dx.plos.org/10.1371/journal.pone.0224854

42. Reed LJ, Muench H. A simple method of estimating fifty per cent endpoints. Am J Epidemiol [Internet]. 1938 May 1 [cited 2024 Aug 12];27(3):493–7. Available from: 10.1093/oxfordjournals.aje.a118408

43. Fobian K, Fabrizio TP, Yoon SW, Hansen MS, Webby RJ, Larsen LE. New reassortant and enzootic european swine influenza viruses transmit efficiently through direct contact in the ferret model. J Gen Virol. 2015;

44. Counsil D argiculture and food. Statistics Pigmeat 2022 [Internet]. 2022. Available from: https://agricultureandfood.dk/prices-and-statistics/annual-statistics

45. Krog JS, Hjulsager CK, Larsen MA, Larsen LE. Triple-reassortant influenza A virus with H3 of human seasonal origin, NA of swine origin, and internal A(H1N1) pandemic 2009 genes is established in Danish pigs. Influenza Other Respi Viruses [Internet]. 2017 May [cited 2018 May 7];11(3):298–303. Available from: http://doi.wiley.com/10.1111/irv.12451

46. Ryt-Hansen P. Surveillance of Influenza A virus in Danish pigs [Internet]. Statens serum institute, Denmark. 2023 [cited 2023 Sep 19]. Available from: https://www.vetssi.dk/overvaagning/overvaagningsprogrammer/overvaagning-af-influenza-a-virus-i-svin-i-danmark

47. Guo H, Rabouw H, Slomp A, Dai M, van der Vegt F, van Lent JWM, et al. Kinetic analysis of the influenza A virus HA/NA balance reveals contribution of NA to virus-receptor binding and NA-dependent rolling on receptor-containing surfaces. PLoS Pathog [Internet]. 2018 Aug 1 [cited 2023 Jan 3];14(8). Available from: https://pubmed.ncbi.nlm.nih.gov/30102740/

48. Du W, de Vries E, van Kuppeveld FJM, Matrosovich M, de Haan CAM. Second sialic acid-binding site of influenza A virus neuraminidase: binding receptors for efficient release. FEBS J [Internet]. 2021 Oct 1 [cited 2023 Jan 3];288(19):5598–612. Available from: https://pubmed.ncbi.nlm.nih.gov/33314755/

49. Hu M, Jones JC, Banoth B, Ojha CR, Crumpton JC, Kercher L, et al. Swine H1N1 Influenza Virus Variants with Enhanced Polymerase Activity and HA Stability Promote Airborne Transmission in Ferrets. J Virol [Internet]. 2022 Apr 13 [cited 2023 Jan 3];96(7). Available from: https://journals.asm.org/journal/jvi

50. Lai JCC, Karunarathna HMTK, Wong HH, Peiris JSM, Nicholls JM. Neuraminidase activity and specificity of influenza A virus are influenced by haemagglutinin-receptor binding. Emerg Microbes Infect [Internet]. 2019 Jan 1 [cited 2023 Jan 3];8(1):327. Available from: /pmc/articles/PMC6455212/

51. Shi Y, Wu Y, Zhang W, Qi J, Gao GF. Enabling the “host jump”: structural determinants of receptor-binding specificity in influenza A viruses. Nat Rev Microbiol 2014 1212 [Internet]. 2014 Nov 10 [cited 2023 Sep 19];12(12):822–31. Available from: https://www.nature.com/articles/nrmicro3362

52. Long JS, Mistry B, Haslam SM, Barclay WS. Host and viral determinants of influenza A virus species specificity. Nat Rev Microbiol 2018 172 [Internet]. 2018 Nov 28 [cited 2023 Sep 19];17(2):67–81. Available from: https://www.nature.com/articles/s41579-018-0115-z

53. Mostafa A, Abdelwhab E, Mettenleiter T, Pleschka S. Zoonotic Potential of Influenza A Viruses: A Comprehensive Overview. Viruses [Internet]. 2018 Sep 13 [cited 2019 Apr 30];10(9):497. Available from: http://www.mdpi.com/1999-4915/10/9/497

54. Mifsud EJ, Kuba M, Barr IG. Innate Immune Responses to Influenza Virus Infections in the Upper Respiratory Tract. Viruses [Internet]. 2021 Oct 1 [cited 2023 Sep 19];13(10). Available from: /pmc/articles/PMC8541359/

55. Thomas MN, Zanella GC, Cowan B, Caceres CJ, Rajao DS, Perez DR, et al. Nucleoprotein reassortment enhanced transmissibility of H3 1990.4.a clade influenza A virus in swine. J Virol [Internet]. 2024 Mar 19 [cited 2024 Jul 15];98(3). Available from: https://journals.asm.org/doi/10.1128/jvi.01703-23

56. Rosário-Ferreira N, Preto AJ, Melo R, Moreira IS, Brito RMM. The Central Role of Non-Structural Protein 1 (NS1) in Influenza Biology and Infection. Int J Mol Sci [Internet]. 2020 Feb 2 [cited 2023 Sep 14];21(4). Available from: https://pubmed.ncbi.nlm.nih.gov/32098424/

57. Ji ZX, Wang XQ, Liu XF. NS1: A Key Protein in the “Game” Between Influenza A Virus and Host in Innate Immunity. Front Cell Infect Microbiol [Internet]. 2021 Jul 13 [cited 2023 Sep 14];11. Available from: /pmc/articles/PMC8315046/

58. Avanthay R, Garcia-Nicolas O, Zimmer G, Summerfield A. NS1 and PA-X of H1N1/09 influenza virus act in a concerted manner to manipulate the innate immune response of porcine respiratory epithelial cells. Front Cell Infect Microbiol. 2023 Jul 26;13:1222805.

59. Thompson WW, Shay DK, Weintraub E, Cox N, Anderson LJ, Fukuda K. Mortality associated with influenza and respiratory syncytial virus in the United States. JAMA [Internet]. 2003 Jan 8 [cited 2024 Oct 24];289(2):179–86. Available from: https://pubmed.ncbi.nlm.nih.gov/12517228/

60. Rothberg MB, Haessler SD. Complications of seasonal and pandemic influenza. Crit Care Med [Internet]. 2010 [cited 2024 Oct 24];38(4 Suppl). Available from: https://pubmed.ncbi.nlm.nih.gov/19935413/

