## Supplementary files 1-10 for "Rapid Surge of Reassortant A(H1N1) Influenza Viruses in Danish Swine and their Zoonotic Potential": SupplementaryFile1.pdf

**Supplementary File 1.** Nucleotide identity (%) between the segments of the inoculum strains selected for the ferret study (A/swine/Denmark/19922-5/2021 and A/swine/Denmark/15063-1/2020) and the zoonotic H1pdm09N1av (A/Denmark/36/2021).

| | A/Swine/Denmark/19922-5/2021 and A/Denmark/36/2021 | A/Swine/Denmark/19922-5/2021 and A/Swine/Denmark/15063-1/2020 |
|---|---|---|
| PB2 | 99.39% (14 nts) | 93.51% (148 nts) |
| PB1 | 99.65% (8 nts) | 94.64% (122 nts) |
| PA | 99.77% (5 nts) | 94.57% (119 nts) |
| HA | 99.00% (17 nts) | 98.00% (34 nts) |
| NP | 99.74% (4 nts) | 94.79% (79 nts) |
| NA | 99.36% (9 nts) | 98.94% (15 nts) |
| M | 99.58% (4 nts) | 96.20% (36 nts) |
| NS | 99.88% (1 nts) | 80.14% (169 nts) |

The number of nucleotide (nt) differences are indicated in brackets.
