## Supplementary files 1-10 for "Rapid Surge of Reassortant A(H1N1) Influenza Viruses in Danish Swine and their Zoonotic Potential": SupplementaryFile4.pdf

**Supplementary File 4.** Sites in the HA protein of H1pdm09 origin being prone to positive selection.

| Pos selected sites | Prob | RBS/AS |
|---|---|---|
| 88S | 0.532 | Cb |
| 145A | 0.779 | RBS |
| 154P | 0.947 | Ca1 |
| 159K | 0.998 | Ca1 |
| 172G | 0.945 | Sa |
| 174S | 0.895 | Sa |
| 178I | 0.552 | Sa |
| 200F | 0.813 | RBS |
| 202I | 0.967 | RBS |
| 203D | 0.99 | Sb |
| 204S | 0.933 | RBS |
| 206R | 1 | Sb |
| 207I | 1 | Sb |
| 338V | 0.971 | - |

Pos selected sites = amino acid position in the HA protein numbered from the first methionine prone to positive selection. Prob = the probability of the site showing positive selection. RBS = receptor binding site. AS = antigenic site of H1 being either Sa, Sb, Ca1, Ca2 and Cb.
