## Supplementary files 1-10 for "Rapid Surge of Reassortant A(H1N1) Influenza Viruses in Danish Swine and their Zoonotic Potential": SupplementaryFile5.pdf

**Supplementary File 5.** H1pdm09N1av clade defining mutations and the prevalence in the H1pdmNx viruses

| Mutation | H1pdmN1av | H1N1pdm/N2sw | RBS/AS |
|---|---|---|---|
| D16N | 63/63 | 11/40 | - |
| R/K60Q | 61/63 | 0/40 | - |
| L/F87I/T | 63/63 | 0/40 | Cb |
| T99A | 60/63 | 0/40 | - |
| S102P | 50/63 | 0/40 | - |
| N/H114D | 62/63 | 0/40 | - |
| Q121H/R | 59/63 | 0/40 | - |
| K/E136N | 63/63 | 3/40 | RBS |
| T137A | 39/63 | 3/40 | RBS |
| N143H | 63/63 | 12/40 | RBS + Sa |
| N/E144D | 63/63 | 10/40 | RBS |
| L145S | 63/63 | 7/40 | RBS |
| K/N/D146S | 63/63 | 0/40 | RBS |
| E/R/N/Q147K | 58/63 | 8/40 | RBS |
| S152A | 63/63 | 10/40 | RBS |
| T/A/G/D172E | 60/63 | 3/40 | RBS + Sa |
| D173N | 61/63 | 15/40 | RBS + Sb |
| N185D | 61/63 | 23/40 | RBS + Ca1 |
| A/T/I202N/S | 62/63 | 0/40 | RBS |
| A/V/G203D | 56/63 | 5/40 | RBS + Sb |
| D/A204S | 63/63 | 1/40 | RBS |
| R/T/S207W | 63/63 | 20/40 | RBS + Sb |
| N211D | 54/63 | 0/40 | RBS + Sb |
| D213N | 60/63 | 2/63 | RBS |
| N/E239D | 61/63 | 11/63 | RBS + Ca2 |
| V267A | 63/63 | 0/40 | RBS |
| A/D278T | 61/63 | 4/40 | - |
| E/D291N | 62/63 | 1/40 | - |
| Q/H319K | 63/63 | 11/40 | - |
| E/D/V/I338T | 62/63 | 0/40 | - |
| K/E/L/T391G | 63/63 | 0/40 | - |

The mutation is defined as the position in the HA protein based on numbering from the first methionine. The letters indicate the amino acid chance at the given position from with the first letter representing the amino acid occurring in the H1N1pdm09/H1pdm09N2sw viruses and the last letter representing the amino acid occurring in the H1pdm09N1av reassortant viruses. Amino acids are named according to the IUPAC nomenclature. The column named RBS/AS indicate if the given mutation is in the receptor binding site (RBS) or in an antigenic site (AS) and which (Sa, Sb, Ca1, Ca2 and Cb). Red = positive selected site.
