## Supplementary files 1-10 for "Rapid Surge of Reassortant A(H1N1) Influenza Viruses in Danish Swine and their Zoonotic Potential": SupplementaryFile7.pdf

**Supplementary File 7.** Overview of the different length and origin of the NS1 protein identified in the H1N1pdm09 and H1pdm09N1av genotypes

| | NS1 length (aa) | H1N1pdm09 Genotype 1 PPPPPP | H1N1pdm09 Genotype 2 PPPPPA | H1pdm09N1av Genotype 1 PPPPPP | H1pdm09N1av Genotype 2 PPPPPA |
|---|---|---|---|---|---|
| *Pdm* | 219 | x | | x | |
| *H1N1av* | 230 | | | | x |
| | 217 | | x | | |

"Pdm" indicates a NS1 origin of H1N1pdm09 whereas "H1N1av" indicates NS1 of H1N1av origin. Aa = indicates Amino acids. There are few exceptions to this generalization of the distribution of the NS1 protein, which is indicated in the phylogenetic tree in Figure 3.
